# Chemical maturation controls bioavailability of Fetuin-A-mineral complexes in biomineralization

**DOI:** 10.1101/2025.01.13.632733

**Authors:** Judith M. Schaart, Luco Rutten, Shaina V. To, Alisha A. Shah, Martijn Martens, Elena Macías-Sánchez, Willi Jahnen-Dechent, Nico Sommerdijk, Anat Akiva

## Abstract

Living organisms must transport calcium and phosphate at high concentrations to enable bone formation without triggering uncontrolled mineral precipitation. Fetuin-A binds calcium phosphate to form soluble calciprotein complexes, but how these complexes contribute to physiological mineralization has remained unclear. Here we show that Fetuin-A-based mineral complexes exist in functionally distinct states that determine mineral bioavailability. Using cryogenic and liquid-phase electron microscopy, biochemical analysis, and osteoblast cell cultures, we demonstrate that small, chemically labile calciprotein monomers directly mineralize collagen fibrils, whereas larger, chemically matured primary calciprotein particles cannot. Instead, these particles require cellular uptake and lysosomal processing to release mineral for matrix deposition. This functional divergence arises from a irreversible chemical transformation of the mineral phase that drives the primary CPP assembly. Together, our findings establish nanoscale chemical maturation as a key control point that separates mineral transport from mineralization, reframing our understanding both physiological bone formation and pathological calcification.

## Introduction

Living organisms face a fundamental challenge in mineral transport: ions such as calcium and phosphate must circulate at high concentrations in blood, but remain soluble to avoid pathological calcification. At the same time, these same ions must be delivered in bulk to support skeletal development and continuous bone remodeling. Still it remains poorly understood how vertebrates solve this paradox - stabilizing mineral during transport while enabling its controlled deposition into bone.

Fetuin-A (FetA) is a liver-produced glycoprotein that is secreted into the blood where it binds calcium phosphate (CaP) to form soluble protein-mineral complexes. Here FetA differs from other proteins e.g. albumin that only bind calcium ions.^1^ This FetA-mineral complexation prevents mineral precipitation and protect soft tissues such as the kidney and blood vessels from ectopic calcification.^1-3^ Yet paradoxically, FetA is also one of the most abundant non-collagenous proteins in bone,^4, 5^ suggesting that beyond its protective role in circulation, it contributes directly to skeletal mineralization.

Protein–mineral complexes formed with FetA exist in multiple states. Small, soluble complexes, called calciprotein monomers (CPMs), have been proposed to carry pre-nucleation mineral clusters, while larger calciprotein particles (CPPs) contain amorphous (primary CPP, Ø50-100 nm) or crystalline (secondary CPP, Ø100-500 nm) calcium phosphate.^2, 4, 6^ CPPs form at increased calcium and phosphate concentrations, first as primary CPPs that upon prolonged ageing transform into secondary CPPs. Both types are strongly associated with chronic kidney disease and vascular pathology.^7-11^ Despite their relevance to physiology and disease, the roles of these different FetA-based complexes in delivering mineral to the extracellular bone matrix have not been elucidated.

Here, we combine cryogenic and liquid-phase electron microscopy with biochemical assays and osteoblast cultures to unravel the formation mechanism of primary CPPs and shed light on their role in the mineralization of collagen in the extracellular matrix (ECM) of bone. We show that the evolution of CPMs into primary CPPs is a two-step process involving a chemical transformation that impacts their collagen mineralization capacity. Small early stage complexes deposit calcium phosphate into collagen fibrils and directly contribute to the mineralization of the bone ECM. In contrast, the larger chemically matured CPPs require cellular uptake and lysosomal processing by the bone-forming osteoblasts before their mineral becomes available for matrix deposition.^12, 13^ These findings imply that beyond capturing excess mineral ions in the blood for clearance through the liver, CPPs may also act as a feedstock supporting bone biomineralization.

## Results

### Formation of Fetuin-A-based protein-mineral complexes

To generate pathological high-phosphate conditions under which the physiological FetA-mineral complexes transform into calciprotein particles, we increased the phosphate concentration of serum-containing cell culture medium at 37 °C (4.75 mM, Supplementary Fig. 1A).^10^ Dynamic light scattering revealed a rapid increase in particle size between 10 and 30 minutes, stabilizing at ∼100 nm, consistent with the complete transition from soluble CPMs to particulate primary CPPs (Fig. 1A). CryoTEM imaging of matured biomimetic CPP solutions showed virtually no CPMs but almost only spherical CPPs ranging from 20 to 100 nm in diameter, alongside protein from the medium (Fig. 1B, Supplementary Fig. 2/3). FetA-mineral complexes and medium proteins were differentiated on the bases of their contrast in the cryoTEM images (Supplementary Fig. 2/3). Selected-area electron diffraction showed diffuse scattering rings characteristic of amorphous calcium phosphate, confirming that the observed spherical complexes were primary CPPs (Fig. 1C). High-resolution images revealed that individual CPPs consisted of 2–5 nm subunits, likely representing aggregated CPMs (Fig. 1B_i_). We note that although FetA has been reported as the main protein supporting CPP formation, the contribution of other serum proteins, e.g. albumin, cannot be ruled out.^14^

**Fig. 1.**
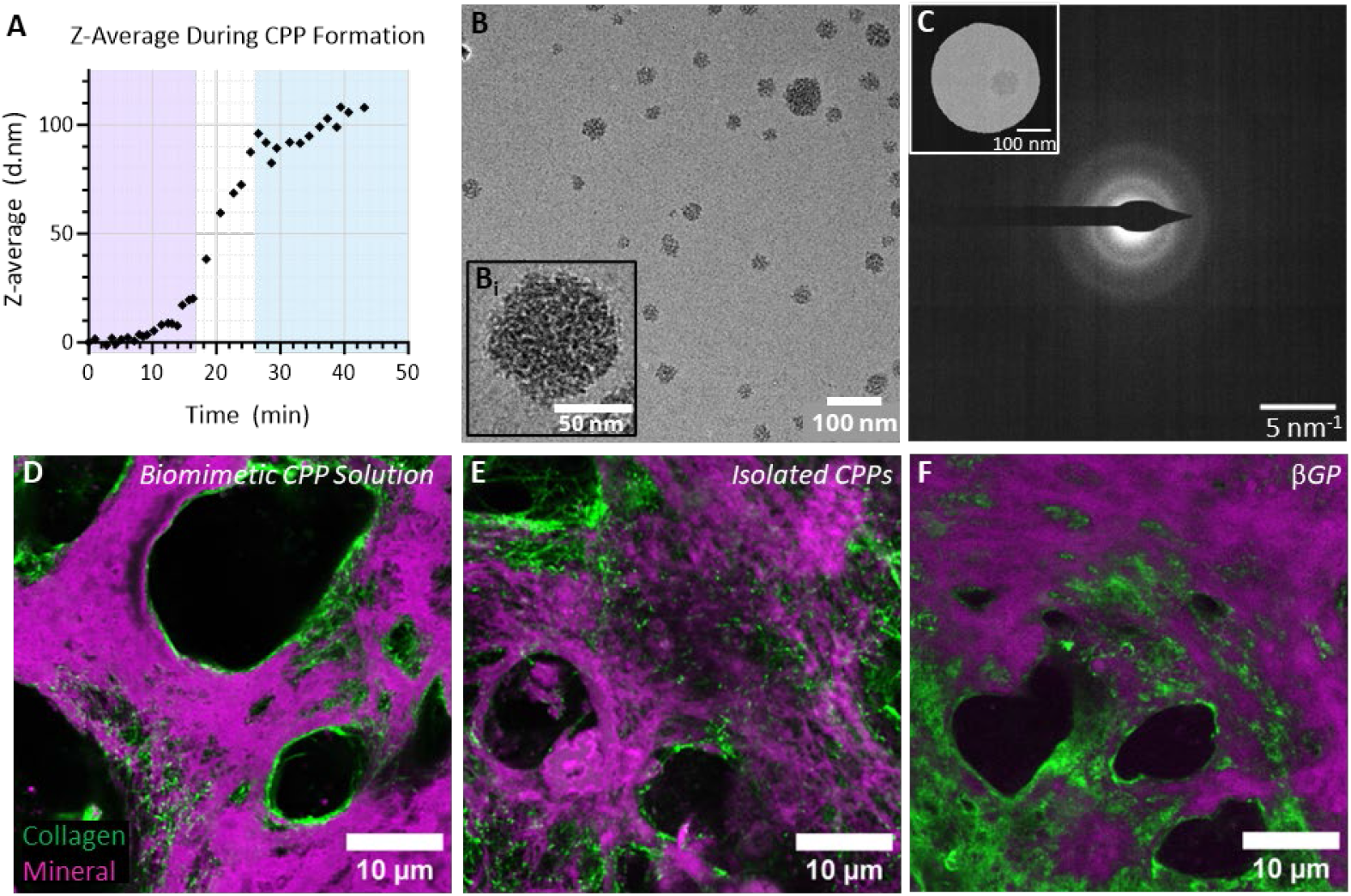
CPP formation and mineralization capacity in osteoblast cultures. (**A**) The transformation of CPMs to primary CPPs in biomimetic CPP solution was monitored by time-resolved DLS. This revealed a stable Z-average signal (close to 0 nm after background correction) in the first 16 minutes after phosphate addition (magenta), followed by a steep increase in Z-average (white), after which a new size equilibrium (∼105 nm) is reached around 25 minutes after the start of the reaction (blue). (**B**) CryoTEM of matured biomimetic CPP solution (24h) shows the presence of spherical complexes (CPPs) with diameters ranging from 20 to 100 nm. (**Bi**) The CPPs are composed of multiple smaller particles, as observed with cryoTEM imaging at higher resolution. (**C**) Diffuse scattering rings observed in selected-area electron diffraction confirms that the observed particles are primary CPPs, carrying amorphous mineral. **D-F**) Fluorescent images of a collagen matrix (CNA35-OG®488, green)) produced by osteoblasts and mineralized (Calcein Blue, magenta) in the presence of (**D**) matured biomimetic CPP solution (containing predominantly CPPs), (**E**) isolated CPPs, or (**F**) β-glycerol phosphate (β-GP). Single channel images are available in Supplementary Fig. 4A. We note lower intensity of the collagen signal in regions with a high mineral signal, due to the presence of the mineral limiting access of the collagen dye.

### CPPs act as mineral source in osteoblast cell cultures

To examine whether these calciprotein complexes could contribute to bone matrix mineralization we exposed matrix-producing mouse osteoblast (MLO-A5) cultures^15^ to matured biomimetic CPP solution as well as to regular cell culture medium supplemented with isolated, preformed CPPs. In both cases, after 14 days of supplementation, fluorescence microscopy revealed matrix mineralization through the overlap of the mineral signal with the labeled collagen fibrils (Fig. 1D-E, Supplementary Fig. 4A). The mineralization by both CPP-containing media was qualitatively similar to that in standard osteogenic cell culture conditions using β-glycerophosphate (βGP) as phosphate source (Fig. 1F), where phosphate is generated by osteoblast-produced alkaline phosphatase (ALP).^15-17^ CryoTEM also identified primary CPPs in the βGP-supplemented culture medium (Supplementary Fig. 1B), underlining that CPP formation in serum is directly linked to increased phosphate concentrations. No mineralization was observed in the absence of added calcium phosphate sources (calciprotein complexes/βGP, Supplementary Fig. 1C).

Cell culture experiments with low [FetA] medium ([FetA]= 0.31mg/mL, Supplementary Fig. 1D-E) using βGP as the mineral source, further illustrated the role of FetA in regulating osteogenic matrix mineralization by controlling CaP precipitation equivalent to the use of crystallization control agents such as poly(aspartic acid).^18^ Here, we observed mineral deposition already after 7 days, compared to 14 days at standard FetA concentration ([FetA]= 2.6 mg/mL, Supplementary Fig. 1D-E), and with poor control over the mineralization process as evidenced by the formation of large globular precipitates, rather than homogeneous mineralization of the collagen matrix (Supplementary Fig. 1F-H). No matrix mineralization was observed when the primary CPPs were added to a cell culture where cellular activity was arrested by chemical fixation. This indicates that primary CPPs can serve as a mineral source during matrix mineralization, but only in the presence of active osteoblasts.

### CPMs Mineralize Collagen, primary CPPs do Not

To further establish the role of osteoblast acivity in CPP-mediated matrix mineralization, we performed cell-free experiments in which we incubated bovine collagen type I fibrils with both freshly prepared and aged biomimetic CPP solutions, containing predominantly CPMs or CPPs, respectively (Supplementary Fig. 1A/3, Supplementary Table 1, Supplementary Notes). CryoTEM revealed intrafibrillar mineralization of the collagen for the CPM solutions (Fig. 2A), in line with previous reports,^18^ but no collagen mineralization was observed with the matured primary CPP-containing solutions (Fig. 2B). Similarly no collagen mineralization was achieved with solutions supplemented with isolated mature primary CPPs, even after prolonged incubation (Fig. 2C). The above findings indicate that under the high-phosphate conditions tested, only CPMs are able to directly deliver mineral to the collagen matrix, whereas CPPs require the involvement of the MLO-A5 cellular activity for their conversion into bioavailable CaP that can mineralize the collagen.

**Fig. 2.**
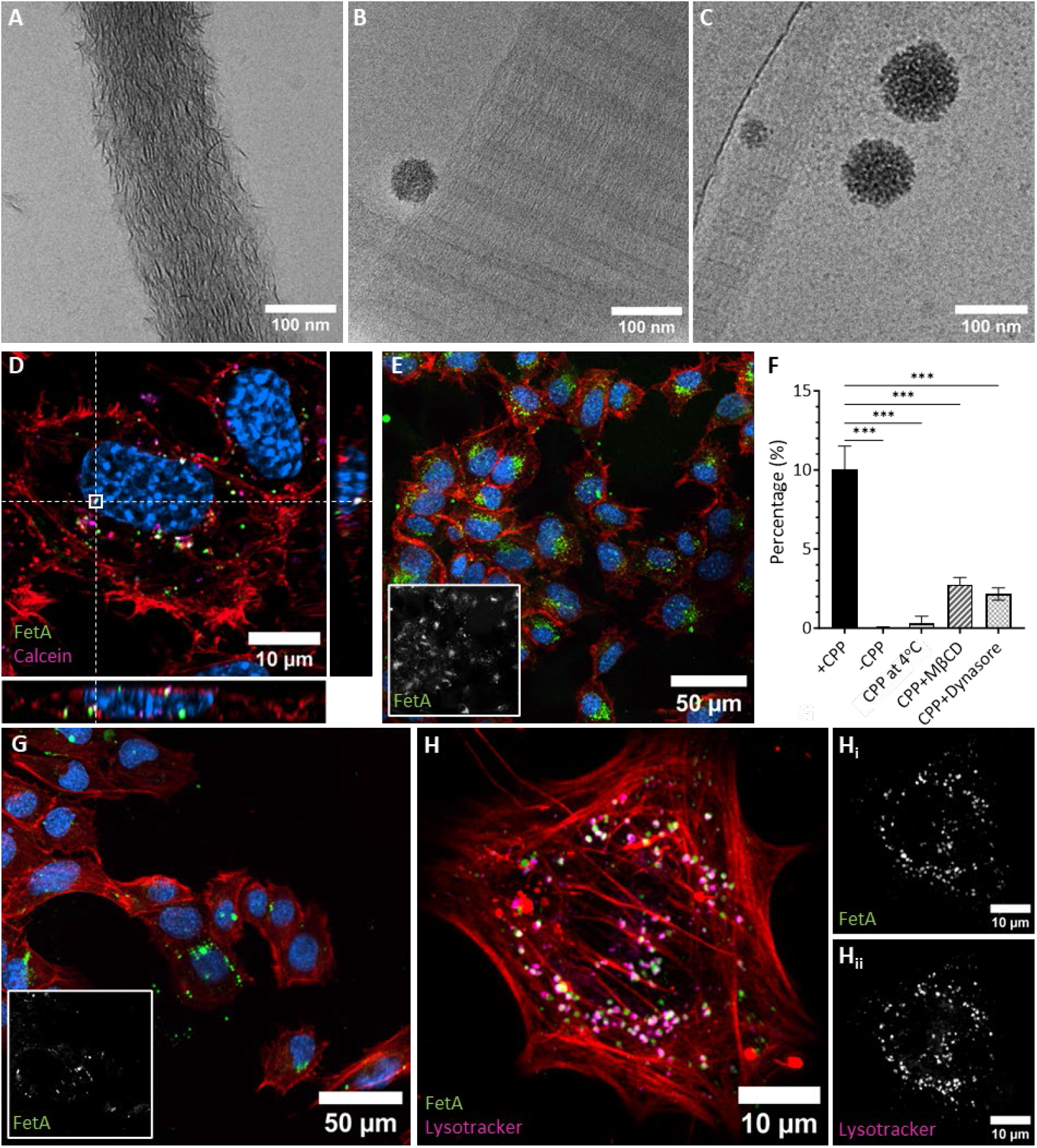
Identification of two mineral delivery pathways by FetA-mineral complexes. (**A-C**) CryoTEM showed (**A**) mineralized bovine collagen fibrils upon incubation (70 h, 37°C) with fresh biomimetic CPM solution See supplementary Fig 3A,C,E), (**B**) no mineralization upon incubation (21 h, 37°C) of bovine collagen fibrils with aged biomimetic CPP solution (24 h incubated after addition of phosphate, See supplementary Fig 3B,D,F), (**C**) no collagen mineralization upon prolonged incubation (70 h, 37°C) of bovine collagen with concentrated, isolated mature CPPs (24 h old). (**D**) Orthogonal views of 3D fluorescence imaging show the presence of double labeled particles (white overlay, FetA-CF®568: green; Calcein: magenta) in the intracellular space, which is bordered by the actin cytoskeleton (red, SiR-actin). Nuclei are indicated in blue (Hoechst 33342). (**E**) Fluorescent imaging of cells (actin: SiR-actin, red; nucleus: Hoechst 33342, blue) cultured in the presence of labeled CPPs (FetA-CF®568, green) showed FetA signal overlapping with cellular area. Inset: Isolated FetA channel. (**F**) Quantification of intracellular FetA signal in the presence and absence of CPPs, and in the presence of CPPs under endocytosis inhibiting conditions (low temperature, MβCD, and Dynasore, respectively). Significance calculated by ANOVA multiple comparison analysis in GraphPad Prism (*** p<0.001). (**G**) Representative images showing MLO-A5 osteoblasts (actin: SiR-actin, red; nuclei: Hoechst 33342, blue) incubated with labeled FetA in the presence of endocytosis inhibitor methyl-β-cyclodextrin. Inset shows the isolated FetA channel. Representative images of the other conditions can be found in Supplementary Fig. 4. (**H**) MLO-A5 osteoblasts (actin: SiR-actin, red) incubated with FetA-CF®568 labeled CPPs (green, **Hi**) and Lysotracker®-DND26 Green (magenta, **Hii**) showing colocalization.

### CPPs are Internalized and Processed via the Endo-Lysosomal Pathway

To understand how osteoblasts process CPPs, we fluorescently labeled both FetA and calcium in the protein-mineral complexes in the biomimetic CPP solution that was added to the MLO-A5 cell cultures (green and magenta, respectivly in Fig 2D). 3D Airyscan microscopy showed dual-labeled CPPs (white in Fig. 2D) within the intracellular space, as outlined by the actin cytoskeleton that indicates the cell borders (red in Fig. 2D). These intracellular CPPs are consistent with active CPP internalization by osteoblasts (Fig. 2D-E). In contrast, no FetA or calcium signal was observed without addition of the CPPs (Supplementary Fig. 4B). Furthermore, the CPP internalization was significantly reduced by the addition of methyl-beta-cyclodextrin^19, 20^ (MβCD, ∼3.6 fold reduction, p=0.0005) or Dynasore^21^ (∼4.6 fold reduction, p=0.0004), two well-known inhibitors of the uptake through cellular engulfment (endocytosis), and completely absent upon incubation at low temperatures (4°C, Fig. 2F-G, Supplementary Fig. 4C-D).

Tracing the intracellular pathway of the fluorescently labeled CPPs by live imaging, using Lysotracker® as a marker for acidic compartments, revealed strong spatial overlap of FetA with lysosomes (Pearson’s coefficient: 0.7),^22^ indicating lysosomal processing after endocytosis (Fig. 2H). Further tracing of the mineral was not possible due to the disintegration of the CPPs into their molecular and ionic components in the acidic environment of the lysosomes. Together, these results imply that bioavailability of the CPP mineral is achieved only after endocytic uptake and acidic lysosomal processing of the complexes by osteoblasts and we infer that this intracellular processing precedes the contribution to matrix mineralization.

### Dynamic assembly of Fetuin-A complexes explains functional differences

To resolve why CPMs but not CPPs directly mineralize collagen, we visualized the CPM-to-CPP transition using liquid-phase electron microscopy (LP-EM). LP-EM allows dynamic real-time visualization of CPP formation with nanoscale resolution, and at the electron dose applied here, without altering the kinetics of the assembly process.^23^ Dynamic imaging of the biomimetic solution during CPP formation within graphene-liquid cells showed the formation of small transient CPPs (tCPPs, diameters ∼5-25 nm) by clustering of CPMs that assembled and disassembled dynamically (Fig. 3A-C, ROIs A1, A2, B1, Supplementary Movie 1-2, Supplementary Fig. 5A). This dynamic process of continuous assembly and disassembly during the CPM-tCPP transition, was further illustrated by color-coding the FetA-mineral assemblies based on their size (Fig. 3A-C, ROIs: A1,A2,B1, Supplementary Movie 1/3).

**Fig. 3.**
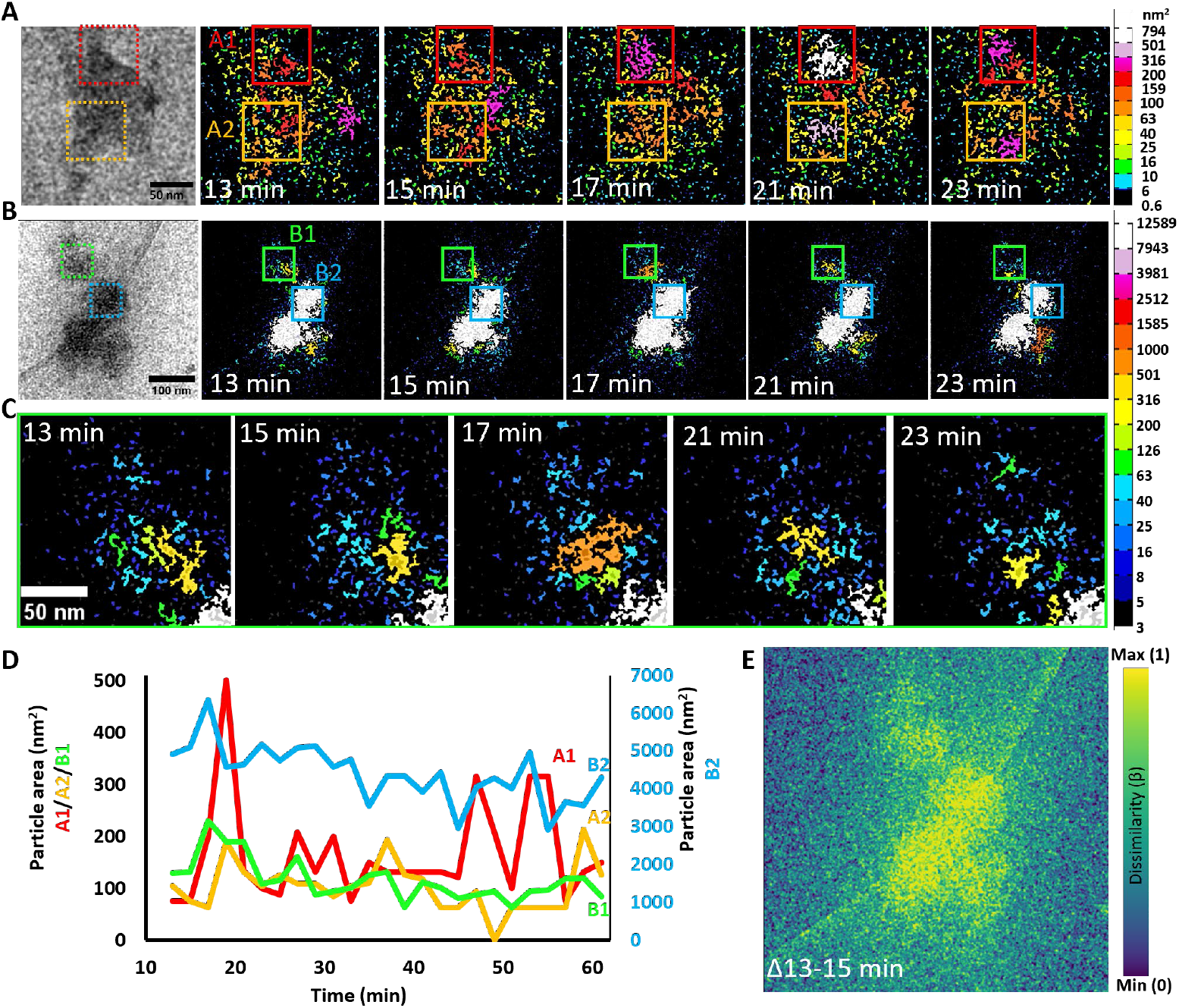
Assembly of stable primary CPPs in phosphate-supplemented cell culture medium is a two-phase process. (**A**) Dynamic LP-EM monitoring of CPP formation in graphene liquid cell loaded with biomimetic CPM solution. Original grey scale TEM image (left), followed by color-based analysis of the particles based on their measured size (projected area, nm^2^) showing the formation of small transient CPPs (<50 nm, tCPP, regions A1 (red) and A2 (yellow)) by assembly and disassembly of CPM clusters. The first image was recorded 13 min after addition of phosphate to the solution. Images were recorded every 2 minutes (Supplementary Movie 1). Scalebar: 50 nm, electron dose: 0.1 e^-^ Å^-2^ per image. (**B**) Time-lapse LP-EM with area-based color-coding (nm^2^) similar as in (**A**) of a graphene liquid cell containing three mature primary CPPs (>50 nm, region B2) and an area in which tCPP (dis)assembly is observed (<50 nm, yellow arrow, region B1). Sequence shows the relative stability of mature CPPs in contrast to the highly dynamic tCPPs. **(C)** Zoom in on the time-lapse of area B1 in **(B)**. Scalebar: 100 nm; electron dose: 0.1 e^-^ Å^-2^ per image. (**D**). Quantification of the particle size per region of interest (as indicated in (**A**) and (**B**)) during 62 minutes (images were recorded every 2 minutes). Note the different y-axis for B2 (right) (**F**) Structural dissimilarity index measure (DSSIM) analysis of sequential frames in the recording of the graphene liquid cell shown in (**B**) showed large changes within the particles, reflecting intraparticle dynamics.

In contrast, the larger primary CPPs (>50 nm) reached a stable size and no longer disassembled, neither did they show an increase in size after reaching a diameter of ∼100 nm, suggesting they had matured to their maximum equilibrium size (Fig. 3B, blue square (ROI B2), Supplementary Movie 2). Internal dynamics were further illustrated by quantifying the size and the number of particles in the regions of interest over time (Fig. 3D, Supplementary Fig. 5C). This confirmed the continuous (dis)assembly of tCPPs, but also indicated fluctuations in the size (projected area) of a mature CPPs. Structural dissimilarity index measure (DSSIM) analysis^24^ further revealed internal dynamics within mature CPPs (Fig. 3E), with the constituting nanoparticulate subunits moving within the CPP perimeter (Supplementary Fig. 5B). The long term stability of graphene-liquid cells allowed prolonged incubation of the CPPs for 14 days. In line with previous reports this led to their transformation into secondary CPPs, as characterized by their oval shape, larger size (∼350 nm long axis), and crystalline mineral content (Supplementary Fig. 5D).

### Chemical maturation drives primary CPP formation

To understand the difference in dynamics and mineralization capacity between early stage and later stage assemblies we analyzed the chemical composition of isolated CPMs and CPPs, as well as the soluble calcium and phosphate concentrations in the solutions. This revealed that CPMs carried mineral complexes with a chemical composition (Ca:P ≈ 0.4, Supplementary Table 1) similar to previously reported pre-nucleation complexes ([Ca(HPO_4_)_3_]^4-^),^25^ while CPPs carried solid hydrated amorphous calcium phosphate (ACP; [Ca_2_(HPO_4_)_3_]^2-^, Ca:P ≈ 0.7) similar to the reported multi-stage development pathway of biomimetic CaP described by Habraken *et al*.^25^. Such chemical difference between early and late stage FetA-mineral complexes could possibly reflect the previously proposed existence of low-density and high-density CPPs.^26^

The chemical transformation associated with this maturation process, explaines why we do not observe a constant equilibrium between the CPMs and CPPs (Fig. 1A,B, Supplementary Figs. 3, 5), but that the system is driven towards the more stable primary CPPs. As expected, the mineral carried by the complexes is in equilibrium with the solution, as indicated by the detected dissolved calcium and phosphate ion concentrations which are in line with the solubility product of ACP.^25^

We propose that the chemical stability of the primary CPPs prevents direct collagen mineralization but enables mineral capture and subsequent relaese through cell-mediated processing. Based on the above analyses we estimate the Ca/protein ratios of the protein-mineral complexes to be 12 ± 2 for the CPMs and 16 ± 4 for the CPPs, respectively (Supplementary Table 1), implying that primary CPPs - formed at higher phosphate concentrations - are more efficient (protein/Ca ratio) in capturing Ca than the CPMs.

### Mineral fate is determined by calciprotein maturation

Our results show that mineral delivery in biology is controlled by the *state* of the protein-mineral complex and not only by the presence of FetA. We demonstrate that small, early-stage FetA-mineral complexes can directly mineralize collagen, whereas larger, chemically matured primary CPPs cannot. Instead, these particles must first be taken up and processed by osteoblasts before their mineral becomes available for matrix deposition. This establishes a functional separation between two mineral transport forms that so far have been treated as variations of the same entity. Calciprotein particles therefore emerge not as inert byproducts of mineral excess, but as a distinct, cell-dependent mineral reservoir. The key implication is that mineral transport does not directly result in mineralization, but that its bioavailability is dependent on the progression of nanoscale chemical CPP maturation and regulated through cellular uptake and processing. This distinction reshapes how we view both normal bone formation and pathological calcification, and clarifies when transported mineral becomes biologically accessible.

## Materials & Methods

### Fetuin-A-mineral complex preparation

The biomimetic CPM/CPP solution applied to cell cultures and collagen fibrils were prepared by supplementing a solution of regular osteoblast cell culture medium (10% FBS (Sigma) in αMEM (Gibco)) with NaH_2_PO_4_ (2.71 µL, 0.5 M stock) and Na_2_HPO_4_ (4.28 µL, 0.5 M stock) to a final volume of 1 mL (Fig. 4). Biomimetic CPM solution was used directly after addition of the phosphate buffers. Biomimetic CPP solution was used after 24h of incubation at 37°C. The final concentrations in the solutions were 2.6 mg/mL Fetuin-A, 1.92 mM Ca^2+^, and 4.75 mM phosphate (Supplementary Fig. 1A). To concentrate the complexes for quantitative analysis, the solutions were spin-filtered using 3 kDa or 300 kDa MWCO Vivaspin 2 filters in a swing bucket rotor (10 min, 4000 g, RT).

**Fig. 4.**
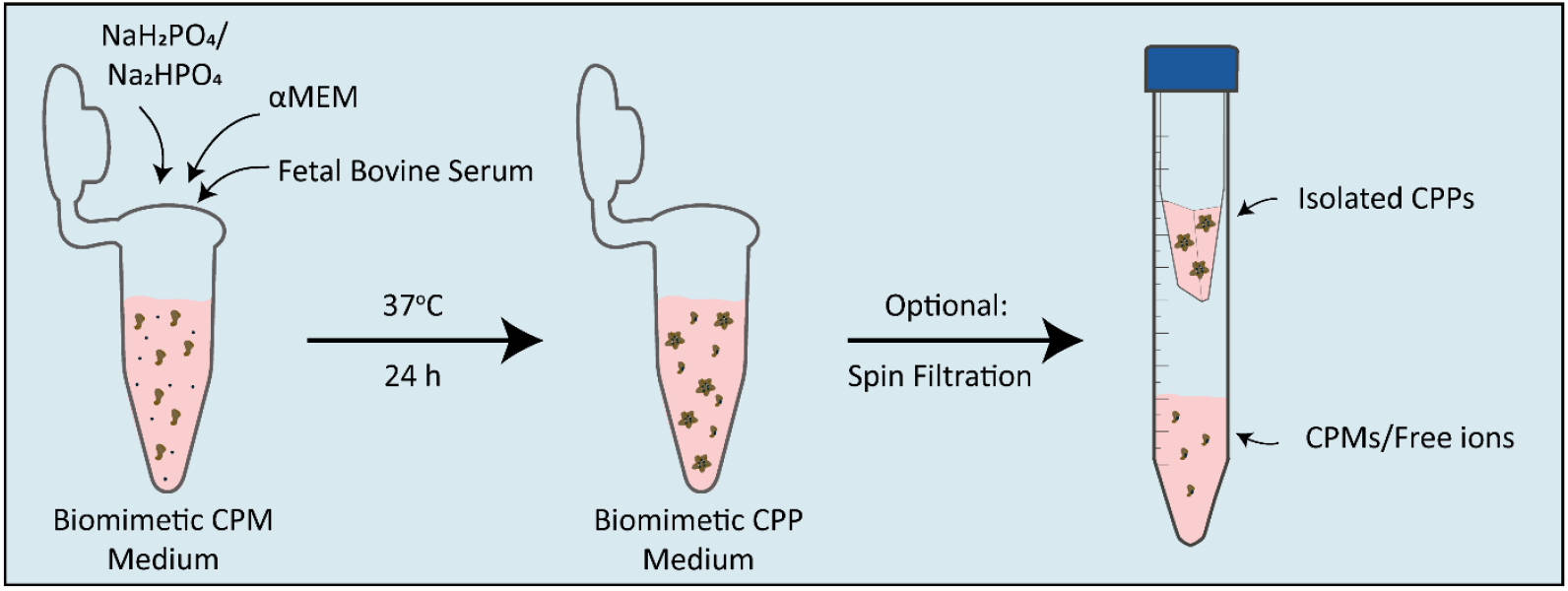
Preparation process of biomimetic solutions containing FetA-mineral complexes in osteoblast cell culture medium. A solution of 10% FBS in αMEM was supplemented with phosphate to form CPM-containing solution. Upon incubation 37°C (24 h) the formation of CPPs was stimulated. If indicated, CPPs were separated from CPMs and free ions by spin filtration.

### DLS

To analyze CPP particle size, DLS measurements were performed on the Zetasizer Nano and the Zetasizer Pro (timelapse). The FetA-mineral complex solution was analyzed directly after production of the particles to determine particle size. For timelapse measurements, 10% FBS in αMEM was analyzed at 37°C before the additional phosphate was added. After phosphate addition, measurements were taken approximately every 90 seconds during the first 40 minutes, while incubating the solution in the Zetasizer at 37°C. Plots were prepared in GraphPad Prism version 10.1.2.

### Western Blot

Western blots were used to analyze the concentration of FetA in FBS and BCS. Three known concentrations of FetA were loaded next to three dilutions of FBS and BCS on a stain-free TGX gradient gel (4-15%, Biorad). After running (200V, 30 min), total protein loading was visualized using a GelDoc (Biorad) and the proteins were transferred to an activated PVDF membrane using a Trans-Blot Turboblotter (Biorad, Mixed MW program). After transfer, the blot was incubated overnight (4°C) in rabbit-anti-FetA (developed at RWTH Aachen) in PBS/0.1% Tween-20 (PBST). Next, the blot was washed 3x in PBST and incubated with a goat-anti-rabbit IRDye-800 secondary antibody, before imaging on an Odyssey Clx (LI-COR). Protein concentrations were quantified based on the net pixel intensity of the bands in FIJI.^27^ The band areas per lane were selected using a consistent ROI frame size based on the largest band area. Next, background areas above and below each band were selected using a consistent ROI frame size. All areas were measured for mean gray values and the net pixel intensity value of the band was derived by subtracting the average of the two background gray values from the band gray value.

### Cell culture

MLO-A5 osteoblasts^15^ were obtained from Kerafast and cultured according to the suppliers instructions. In short, cells were proliferated in αMEM supplemented with 5% FBS and 5% BCS (Hyclone, SH30073) in a humidified incubator (37°C, 5% CO_2_) and split 1:20 twice a week after detaching with Trypsin/EDTA (0.05%/0.53 mM).

For mineralization experiments, MLO-A5 cells were seeded on HCl (1 M) washed and ethanol (70%) sterilized coverslips (#1.5). The cells were proliferated until confluent before osteogenic medium was applied. For classical mineralization, the medium consisted of 10% FBS in αMEM, supplemented with 100 µg/mL ascorbic acid (AA), 100 U/mL-100 µg/mL Penicillin/Streptomycin (PenStrep), and 4 mM of beta-glycerophospate (βGP). For mineralization in the biomimetic CPP solution, the CPP preparation was removed from the water bath after 24h of incubation and supplemented with AA (100 µg/mL) and PenStrep (100 U/mL-100 µg/mL). For mineralization in the presence of a low concentration of FetA, the 10% FBS in the medium was replaced by 10% BCS. In all mineralization experiments, the medium was refreshed three times per week.

Collagen production and mineralization were monitored at fixed time points (d7/d14) by fixing the cells in paraformaldehyde (PFA, 4%, RT, 15 min) and staining with CNA35-OG®488^28, 29^ (1 μM) and Calcein Blue (0.1 mg/mL) in PBS (1h, RT). After staining, the samples were washed twice in PBS and mounted in Vectashield (non-hardening).

Fluorescence images were recorded on a LSM900 upright with Airyscan® 2 detector (Carl Zeiss Microscopy GmbH, Germany), using the 405 nm and 488 nm lasers. Overviews were recorded using a 20x (NA 0.8) objective in SR-4Y mode. High resolution z-stacks were recorded using a 63x oil immersion objective (NA 1.4), using SR Airyscan mode.

### CPP preparation and isolation for cell culture in low [FetA] medium

To confirm that CPPs are a functional mineral source for osteoblasts, we added isolated, preformed CPPs to medium with low FetA concentration. For isolation and dilution in cell culture medium, complexes were prepared by mixing pre-warmed FBS (400 µL) with NaCl (100 µL, 140 mM), phosphate buffer (250 µL, 40 mM NaCl, 19.44 mM Na_2_HPO_4_, 4.56 mM NaH_2_PO_4_·1H_2_O, pH 7.4) and calcium stock solution (250 µL, 140 mM NaCl, 40 mM CaCl_2_·2H_2_O, 100 mM HEPES). The mixture was incubated at 37°C (10 min). Particles were pelleted (20000 g, 20 min, 4°C) and pellets were resuspended in αMEM. Then, CPPs were isolated and further concentrated on 300 kDa MWCO Vivaspin 2 filters (10 min, 3000 g, RT), for use in cell culture experiments.

To prevent overloading of the culture with FetA, which is also naturally present in the complete culture medium because of FBS supplementation, the fetal calf serum (FBS) in these samples was replaced by bovine calf serum (BCS). This serum contained approximately 8.5x less FetA ([FetA_BCS_] = 3.1 mg/mL; [FetA_FBS_] = 26.4 mg/mL) as determined by western blotting (Supplementary Fig. 1B). The final medium used for these experiments consisted of 10% BCS in αMEM, supplemented with 100 µg/mL AA, PenStrep (100 U/mL-100 µg/mL) and isolated CPPs providing an additional 1.6 mM calcium, as determined based on the calcium concentration of the concentrated CPP solution. The suitable concentration of CPPs for cell culture experiments was determined empirically, by titrating the mineralization (1.6 mM/3.2 mM/6.4 mM Ca^2+^supplementation by CPPs) and analyzing the resulting cultures with fluorescent stainings for collagen en mineral. From this it was concluded that 1.6 mM Ca^2+^ supplementation by CPPs was sufficient.

#### Calcium assay isolated CPPs for cell culture

The volume of isolated CPPs to be added to cell culture medium to reach a concentration of 1.6mM Ca^2+^ supplementation was determined for each CPP preparation. The calcium concentration of the isolated CPPs in solution was determined using an O-cresolphthalein-based assay (Randox). Calcium was dissolved from the CPPs by mixing two volumes of CPPs sequentially with one volume of HCl (0.6 M) and one volume of 40 mM NH_4_Cl/5% NH_4_OH. Samples were diluted 125x and 250x in dH_2_O and were mixed with the prepared kit reagents. The kit reagents (component A and B) were mixed in equal volumes before use and 250 µL was added to the samples and CaCl_2_ standards (20 µL). After incubation (15 min, RT), the absorption was measured with a platereader (570 nm, Biorad Benchmark Plus).

### Cell-free mineralization

For cell-free mineralization experiments, 5 µL of bovine tendon collagen (0.8 mg/mL collagen in 0.5 mM acetic acid) was loaded on a gold 200 mesh plasma treated Quantifoil R2/2 grid. After 10 min, the excess liquid was blotted away. The collagen-loaded grid was incubated for 21 h or 70 h on a droplet of biomimetic CPP solution (either directly after preparing (24 h), or after concentrating CPPs 6.25x on a Vivaspin 300 kDa MWCO filter (70 h)) in 100% humidity at 37°C. Next, the grids were vitrified using a VitroBot Mark IV and images were acquired using a JEOL JEM 2100 (Jeol Ltd., Tokyo, Japan) equipped with a Gatan 890 Ultrascan (Pleasanton, CA, USA) and LaB6 filament operated at 200 kV using low dose software (Serial EM).

To study the mineralization capacity of the biomimetic solution before CPP formation, collagen-loaded TEM grids were incubated in a biomimetic CPM medium, directly after the addition of phosphate to the solution. Grids were incubated for 70 h in 100% humidity before vitrification and imaging.

### CryoTEM

For cryoTEM, FetA-mineral complexes in solution were vitrified using a VitroBot Mark IV and images were acquired using a TALOS F200C-G2 (ThermoFisher, Eindhoven) equipped with a Falcon4i direct electron detector, operated at 200 kV. To analyze the presence of CPP in βGP-supplemented cell culture medium, a sample of medium that had been incubated on a differentiating culture was taken at day 7 of differentiation before medium exchange and vitrified using as described above, before imaging at the JEOL JEM 2100 with a Gatan 890 Ultrascan (Pleasanton, CA, USA).

### Fluorescent Fetuin-A/mineral complex preparation

NaHCO_3_ buffer (1 M, pH 8.27 at RT) was prepared. A small amount of CF®568 succinimidyl ester (Merck) was dissolved in DMSO (8 μL) and the dye concentration was determined based on absorbance (Nanodrop 2000c, ε=100.000, CF=0.08, Abs_max_=562 nm). The initial concentration of the FetA monomer solution (purified on a Sephadex column) was determined based on the absorbance measured with the Nanodrop (E_1%_ of FetA: 0.45 mg ml^-1^ mm^-1^ at 280 nm; 23.8 mg/mL). The protein was diluted in PBS to reach a final concentration of 5 mg/mL, then NaHCO_3_ buffer was added to the protein with final concentration of 0.1 M NaHCO_3_ and the dye was added with a 1:10 molar ratio (protein:dye; final reaction mixture: 140.5 μL of 5 mg/mL FetA, 14.1 μL of NaHCO_3_ buffer, 7 μL of 15.7 mM CF568). The protein-dye solution was vortexed briefly and incubated for 1 h at room temperature in the dark on a shaker (300 rpm). Next, the solution was purified twice with an equilibrated Zeba Spin 7K MWCO Desalting column (ThermoFisher) via centrifugation (1000 g, 2 min) to remove the free dye. The concentration of labeled FetA (∼5.3 mg/mL) and the degree of labeling (DOL) of FetA were determined to be around 3, based on measurements of the absorbance (562 nm (fluorescent dye), 280 nm (protein concentration)) and calculated using the following equations.

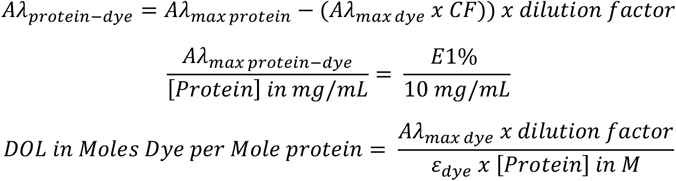

For fluorescent labeling of the FetA-mineral complexes, labeled FetA was added in a molar ratio of 1:100 before the incubation was started. Additionally, for dual labeling, Calcein Green (1 µg/mL in 100 mM KOH) was added during the last hour of incubation.

### Endocytosis inhibition assay

MLO-A5 cells (1000 cells per cm^2^) were seeded on coverslips (#1.5) and cultured for 3 days. Biomimetic CPP solution was prepared as described above, in the presence of fluorescently tagged FetA and Calcein Green. The cells were incubated with the biomimetic medium supplemented with AA (100 µg/mL) and PenStrep (100 U/mL-100 µg/mL) for 4h under variable conditions: standard conditions, at low temperature (4°C), and in the presence of methyl-beta-cyclodextrin (2.5 mM) or Dynasore (40 μM). For endocytosis inhibiting conditions (low temperature/methyl-beta-cyclodextrin (2.5 mM)/Dynasore (40 μM)), the cells were pre-incubated for 30 mins in the inhibitory condition before the proliferation medium was exchanged for biomimetic CPP solution, supplemented with AA and PenStrep. After incubation, the cells were washed with PBS (3x), fixed in 4% PFA (15 min, RT), and stained with Hoechst 33342 (1 μg/mL) and silicon-rhodamine F-Actin (SiR-Actin, Spirochrome, 1 μM) for 1 h. Next, the coverslips were washed twice in PBS and mounted with Vectashield on glass slides.

Fluorescence images were recorded on a LSM900 upright with Airyscan 2 detector. Overviews were recorded using a 20x Plan-Apochromat objective (NA 0.8) in SR-4Y mode. High resolution z-stacks were recorded using a 63x Plan-Apochromat oil immersion objective (NA 1.4), using SR Airyscan mode. For quantification of the intracellular FetA signal, 20x images were processed in FIJI.^27^ Based on the actin signal, a mask for cellular localization was made using automated thresholding (Li ^30^) and holes in the cell mask were filled to cover the full cellular surface. The cellular surface area was calculated by measuring the area of the actin mask using the Analyze Particles function (size: 5 or 20-infinity (5 positive and negative control, 20 for experimental conditions)) and verified manually using the identified boundary boxes. Next, the particles overlapping with the cellular area were selected using the AND function on the actin mask and the FetA channel. The FetA channel was automatically tresholded (Yen ^31^) to prepare a mask and particle analysis (size: 0-infinity, circularity: 0-1) was performed to measure the sum of the surface area calculated with the ROI manager. The calculated surface values were used to determine the percentage of the cellular area that overlapped with the FetA signal. Statistical testing was performed in GraphPad Prism version 10.1.2 (GraphPad Software, Boston, Massachusetts USA, www.graphpad.com) by performing a one-way ANOVA followed by Dunnett’s multiple comparisons test.

### Sub-cellular CPP localization

MLO-A5 cells were seeded and cultured as described for the endocytosis inhibition assay. After 3 days in proliferation medium, the cells were incubated with biomimetic CPP solution containing FetA- labeled protein-mineral complexes for 4 to 6 h and stained live with Hoechst 33342 (1 μg/mL) and SiR-actin (1 μM) in the presence of Verapamil (10 μM) for 1 h. Shortly before imaging, the biomimetic CPP solution was replaced by imaging medium without FetA-mineral complexes, consisting of 10% FBS in αMEM, supplemented with 100 µg/mL AA, PenStrep (100 U/mL-100 µg/mL), Verapamil and Lysotracker® Green DND-26 (0.5 μM). The cells were imaged live using an LSM900 upright with Airyscan 2 detector in SR-4Y mode, using a 63x water dipping objective (W Plan-Apochromat, NA 1.0).

### Liquid-Phase Electron Microscopy

Copper chips coated with graphene by CVD were placed inside a Naiad-MOD-1 (VitroTEM) and the copper was etched away with 0.2 M ammonium persulfate. Next, the free-floating graphene was washed with Milli-Q. A small volume (40 µL) of biomimetic CPM solution was prepared in cell culture medium as described above (αMEM/10% FBS/NaH_2_PO_4_ (0.11 µL, 0.5 M stock)/Na_2_HPO_4_ (0.17 µL, 0.5 M stock)). 5 µL of the mixed solution was applied on a gold 200 mesh ultrathin carbon film finder TEM grid (CF200F1-AU-UL, Electron Microscopy Sciences) and samples were encapsulated in 96% humidity with graphene by automated loop-transfer using the Naiad-MOD-1 system (VitroTEM).

After encapsulation, the grid was transferred to an Elsa cryo-transfer holder (Gatan) and placed in a TALOS F200c G2 TEM (Thermo Fisher) equipped with a Falcon 4i direct electron detector (Thermo Fisher) operated at 200 kV. Images were acquired every 2 minutes at 37 °C during CPP formation, using low-dose imaging (0.1 e^-^ Å^-2^) until the samples had been incubated 1 h. Imaging started 13 min after the addition of phosphate. After 1 h of low-dose imaging, a high-contrast snapshot was recorded (10 e^-^ Å^-2^).

### Chemical analysis of FetA-mineral complexes

Concentrations of calcium, phosphate and protein in the fractions of the biomimetic solution were determined based on standardized tests from the hospital (Diagnostics Laboratory Radboudumc). FetA-mineral complexes were prepared in 10% FBS in αMEM (1 mL), as described above, and isolated directly after phosphate addition (t=0, biomimetic CPM solution) or after 24h of maturation (t=24. biomimetic CPP solution), using spin filtering with 300 kDa MWCO Vivaspin filters (4000 g, 10-15 min, 4 °C; CPP in retentate, CPM and ions in flowthrough) followed by filtering of the flowthrough with 3 kDa MWCO Vivaspin filters (4000 g, 10-15 min, 4 °C; CPM in retentate, ions in flowthrough). The separated CPP and CPM fractions were diluted to a volume of 250 µL in HEPES (50 mM, pH 7.4). Next, all samples were submitted for determination of the calcium, phosphate and protein concentrations. Calcium measurements were based on a photometric test with 5-nitro-5’-methyl-BAPTA and EDTA. Phosphate concentration was determined based on a photometric test with ammonium molybdate. Protein concentration was determined using a colorimetric assay in which a biuret complex with Cu^2+^ was formed in an alkaline solution. Photometric and colorimetric measurements were performed on a ROCHE Cobas 8000, module C702.

### Image Processing

LP-EM images were processed with SenseAI by applying a 2D gaussian blur (x:1, y:1, z:2) and training a dictionary (size 36, 13×13 instance, batch size 32000, sparsity limit 12, learning rate 0.85). Next, the images were imported in Dragonfly 3D World 2024.1 (Comet Technologies Canada Inc., Montreal, Canada; software available at https://www.theobjects.com/dragonfly) and slice registration was performed to align the images. First, manual adjustments were made to remove large jumps. Then automated registration was performed using “optical flow” and allowing only translation. Lastly, manual adjustments were made to optimize the alignment. After aligning, liquid pockets were cropped out of the image and FIJI (v1.53t)^27^ was used to apply thresholding for particle analysis with default settings (size: 0-infinity, circularity: 0.00-1.00) to determine the size of the particles. Next, the ROI color coder from the Broadly Applicable Routines (BAR) plugin^32^ in FIJI was used to color the particles based on their size (logarithmic scale bar). Additionally, cropped liquid pockets (without SenseAI processing) were analyzed by DSSIM analysis in FIJI (standard deviation 1.5, size 11, per-frame contrast, α=0 β=1 γ=0).^24^

To differentiate between proteins in solution and CPMs, the cryoEM images were processed in FIJI. The images were averaged (5 pixels) and a threshold of 99.75 was applied to create a binary image that showed the CPMs and CPPs, but not the other proteins in the solution.

Fluorescence images were processed in FIJI. For visualization purposes, window/level values were adjusted. In cases of comparison between multiple conditions, these settings were adapted per experiment for all conditions. Z-stack maximum intensity projections were prepared for visualization and stacks were resliced (left and top) to prepare orthogonal views.

## Acknowledgements

The authors would like to thank Steffen Gräber from RWTH Aachen University Hospital for the purification of the Fetuin-A used for fluorescent labeling and FBS/medium supplementation. The authors also thank Thomas Kock and Gregory Schneider from Leiden University for supplying the graphene coated chips used in the LP-EM experiments. We thank Joost Hoenderop and Jeroen de Baaij from Radboudumc Nijmegen for their critical evaluation of the manuscript..

## Author contributions

Conceptualization: JS, LR, EMS, NS, AA

Methodology: JS, LR, ST, AS

Investigation: JS, LR, ST, AS, MM

Visualization: JS, LR, AA

Funding acquisition: EMS, WJD, NS, AA

Project administration: NS, AA

Supervision: JS, NS, AA

Writing – original draft: JS, NS, AA

Writing – review & editing: JS, LR, ST, AA, EMS, WJD, NS, AA

## Data and materials availability

All data needed to evaluate the conclusions in the paper are present in the paper and/or the Supplementary Materials. The original data files will be made available in the Radboud Data Repository with DOI 10.34973/m4fq-6146 (upload in progress, link not publicly available yet).

