## Supplementary Materials for "Chemical maturation controls bioavailability of Fetuin-A-mineral complexes in biomineralization"

**The PDF file includes:**

Notes
Supplementary Figures 1 to 5

**Other Supplementary Materials for this manuscript include the following:**

Supplementary Movies 1 to 3
Supplementary Table 1

**Notes**

Notes on the composition analysis of the FetA-mineral complexes

The composition of the different fractions of FetA-mineral complexes (CPP/CPM/Ions) were determined using standardized hospital blood testing. It must be noted that here, the total protein concentration is measured, which also includes additional serum proteins like albumin. To determine the concentration of FetA, we therefore base our calculations of the FetA concentration on the calculated concentration of FetA in FBS, based on western blotting. Throughout all experiments, the same lot of FBS and BCS was used. Therefore, minimal interexperimental variability is expected for the FetA concentration.

Differentiate protein, CPM and CPP

To differentiate between proteins in solution and CPMs, cryoTEM images of freshly made biomimetic CPP solution filtered over a MWCO 300 kDa filter (Supplementary Fig. 2A) and 24 h incubated CPP filtered over a MWCO 300 kDa filter (Supplementary Fig. 2D) were acquired. All images, including the CPP image (Supplementary Fig. 2E), were tresholded using one fixed value based on the CPM image (Supplementary Fig. 2B). The overlays (Supplementary Fig. 2C, 2F) show the differentiation between medium proteins and CPMs.

Validity of LP-EM experiments

In addition to the stable large CPPs observed in Fig. 3B, we also monitored a small tCPP observed alongside the mature CPPs in the same liquid pocket (Fig. 3B, region B1 (green)). This tCPP still showed similar growth/disassembly dynamics (Fig. 3C, Movie S3) as the previously discussed tCPPs (Fig. 3A). This suggests that while feedstock (CPMs) was still present in the solution that allowed for growth of tCPPs, larger matured CPPs had been stabilized in size and shape by internal interactions.

Additionally, secondary CPP formation (Supplementary Fig. 5E) indicates that the CPPs in the liquid pocket undergo similar development as previously observed in bulk samples[1] and that the confined environment of the liquid pockets does not restrict the CPP dynamics that are responsible for maturation or growth.

### Supplementary Figures

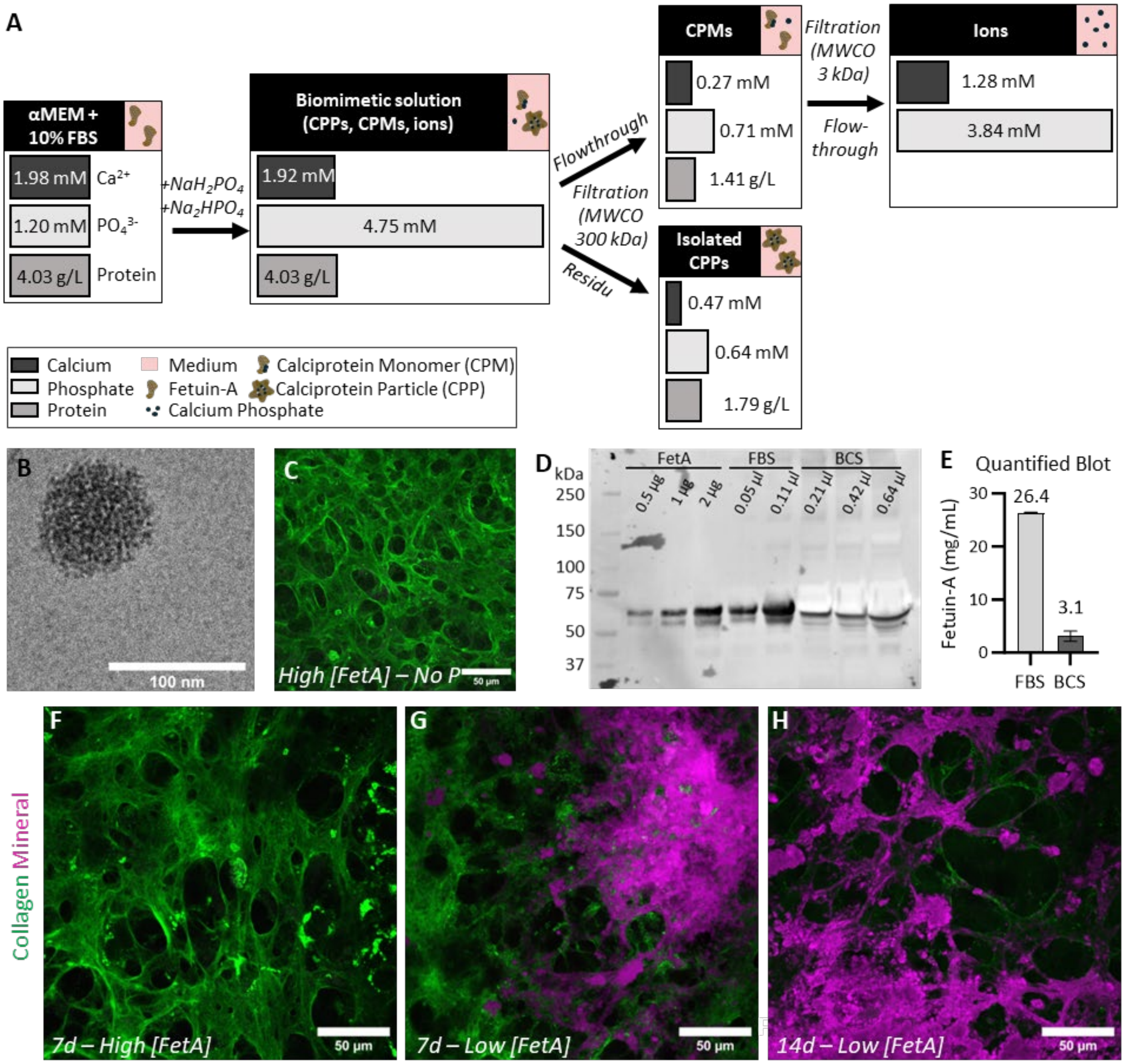


**Supplementary Figure 1.**

(**A**) Characterization of the biomimetic mixture and the separate components, as determined by standardized hospital diagnostic blood-testing to quantify protein, calcium and phosphate content. (**B**) CryoTEM of FBS-supplemented (high [FetA]) medium (αMEM) from matrix-producing cultures (d7) supplemented with βGP shows the presence of CPPs. (**C**) Staining of collagen (CNA-OG®488, green) and mineral (calcein blue, magenta) after 14 days of differentiation in the absence of a mineral source showed no mineralization of the produced collagen matrix. Scalebar: 50 µm. (**D**) Quantification of FetA concentration in FBS and BCS by western blotting. FBS, BCS, and FetA standards were loaded on a stain-free TGX gradient gel (4-15%) and transferred to a PVDF membrane. Immunodecoration against FetA was used to visualize the specific protein bands. (**E**) Quantification of the western blot (**D**) was performed in FIJI software, based on band intensities. (**F-H**) Fluorescent staining of collagen (CNA-OG®488, green) and mineral (calcein blue, magenta) of matrix producing MLO-A5 osteoblast cultures (scalebars: 50 µm): (**F**) In medium with high [FetA], containing 10% FBS, no mineral deposition is observed at d7 of matrix production. (**G**) In medium with low [FetA], containing 10% BCS, mineral deposition is observed at d7 of matrix production. (**H**) In medium with low [FetA], containing 10% BCS over-mineralization is observed at d14 of matrix production.

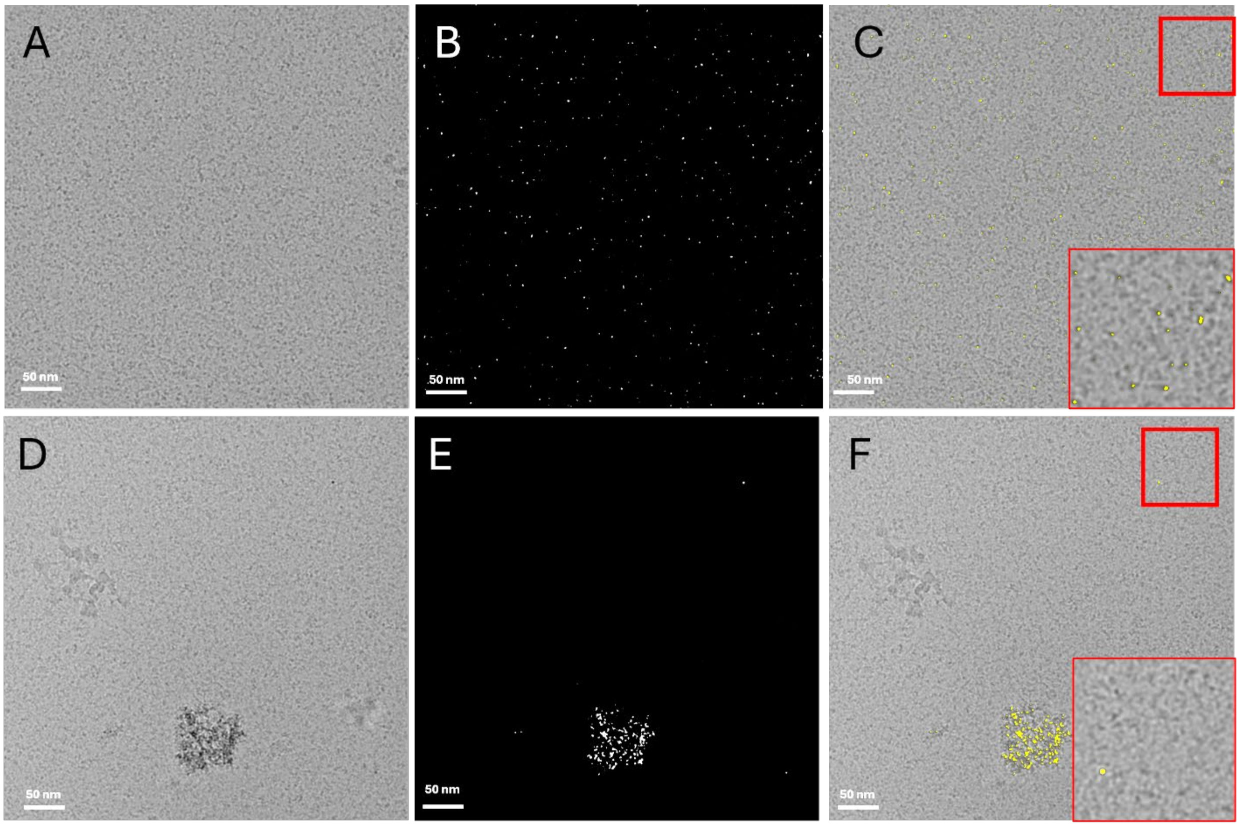


**Supplementary Figure 2. Validation of the capability to distinguish between CPMs and other proteins in solution in cryoTEM images.** (**A**) CryoTEM image of isolated CPMs (MWCO 300 kDa spin filter) and (**B**) corresponding thresholded image mask identifies the CPMs and other proteins by contrast. (**C**) Overlay of the cryoTEM image and the tresholded mask. (**D**) CryoTEM image of isolated CPP solution and (**E**) corresponding thresholded masks show predominantly CPPs and only a few CPMs. (**F**) Overlay of the cryoTEM image and the mask.

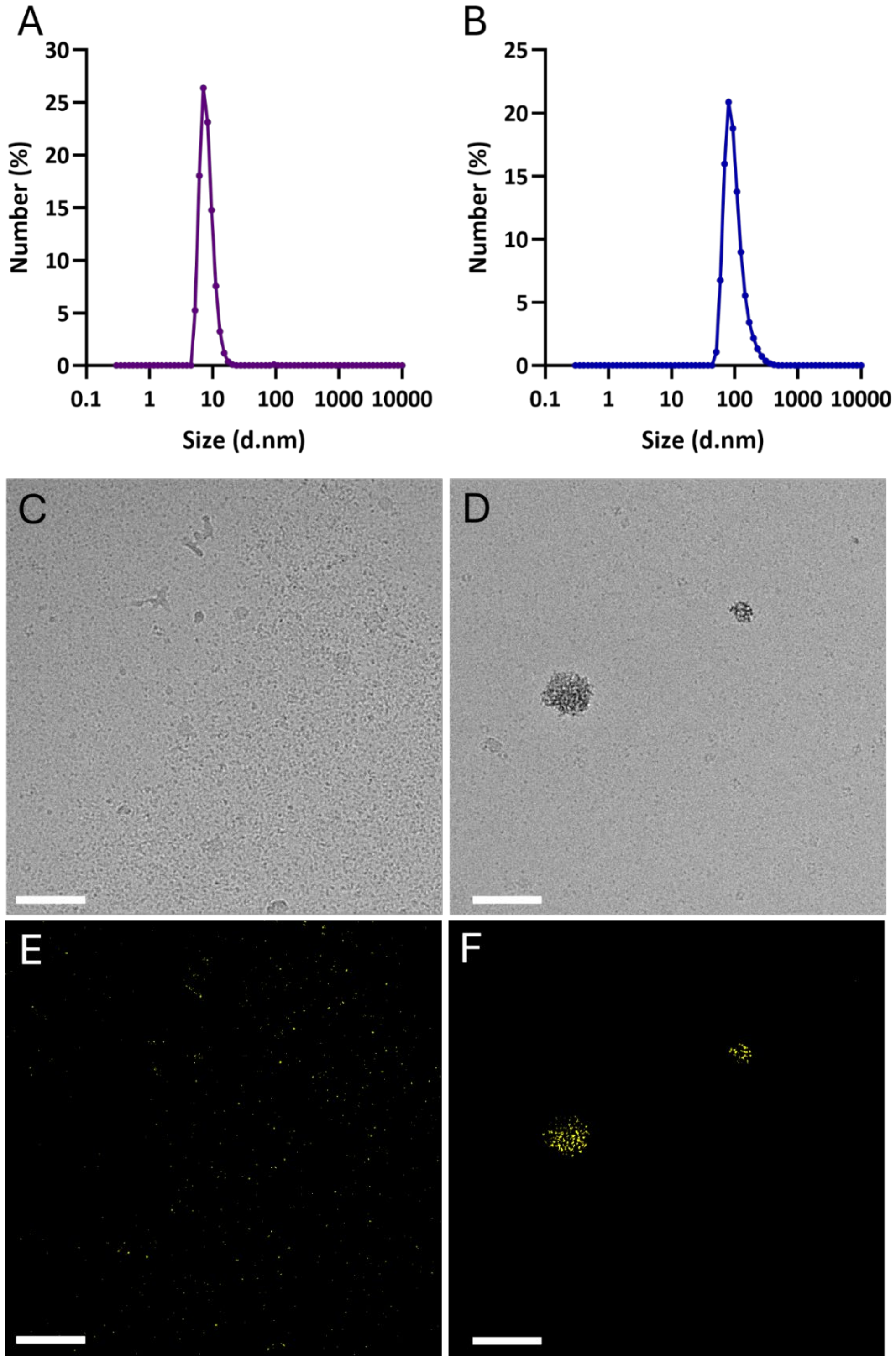


**Supplementary Figure 3. Characterization of CPM and CPP in biomimetic mineralization solution show exclusively CPM at t=0h and predominantly CPP at t=24h.** (**A**) DLS number plot recorded directly after preparation of the biomimetic solution (t=0h). (**B**) DLS number plot recorded after 24h of incubation at 37$^{o}$C. (**C**) CryoTEM of the biomimetic solution at t=0 shows the presence of CPMs, but no CPPs. (**D**) CryoTEM after 24h of incubation at 37$^{o}$C shows the presence of primary CPPs. **E,F**) Threshold masks to distinguish CPM and medium proteins in images (**C**) and (**D**) respectively. Scale bar: 100 nm.

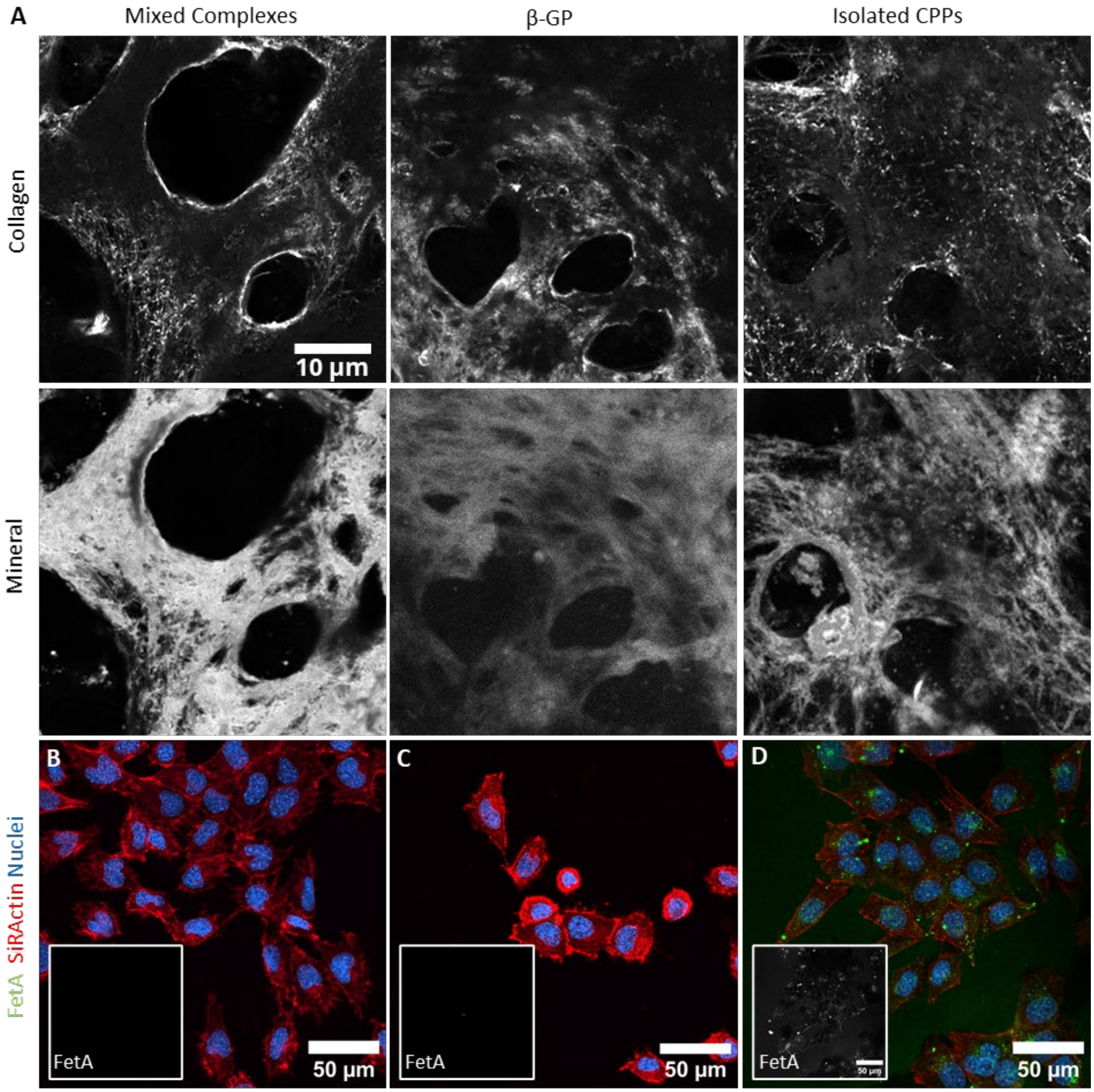


**Supplementary Figure 4.**
(**A**) Single channel images of Fig. 1F-H show overlapping patterns in collagen (top) and mineral (bottom) signal and fading of the collagen signal upon high mineralization, due to low accessibility of the dye to the collagen. (**B-D**) Representative images of cells incubated with (**B**) no CPPs, (**C**) FetA-labeled CPPs at 4°C, (**D**) FetA-labeled CPPs in the presence of endocytosis inhibitor Dynasore. FetA is shown in green, while the cells are indicated by actin (red) and nuclei (blue). Insets show single channel images of the FetA channel.

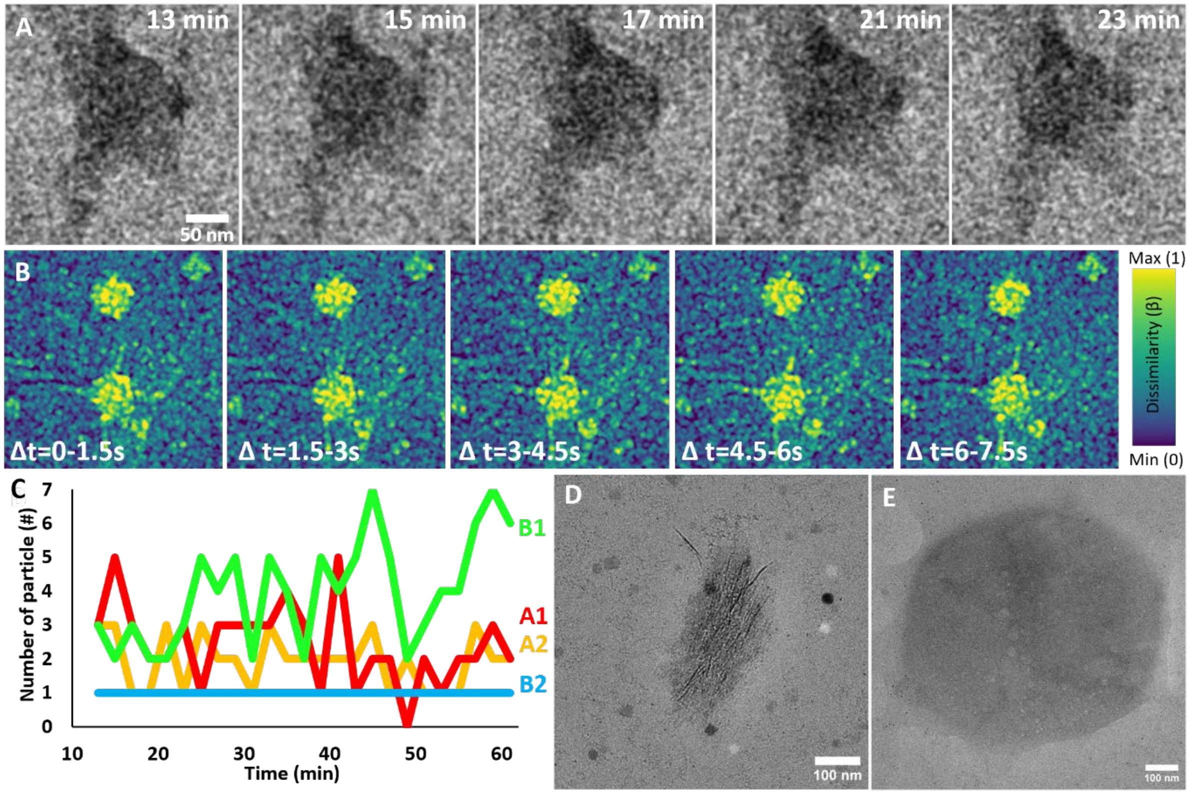


**Supplementary Figure 5.**
(**A**) Timelapse LP-EM images of the graphene-liquid cell shown in Fig. 3A. The first image was recorded 13 min after addition of phosphate to form the biomimetic solution. Images were recorded every 2 minutes (0.1 e- Å-2 per image) and processing with SenseAI was performed as described in the methods. (**B**) DSSIM analysis of sequential frames, recorded with a high time resolution (1.5 s between frames (as shown by Rutten *et al.*[2])) shows intraparticle dynamics within mature CPPs. (**C**) Quantification of the number of particles per region of interest (as indicated in Fig. 3A and Fig. 3B) during 62 minutes (images were recorded every 2 minutes). (**D**) Secondary CPP in a graphene liquid cell, recorded after 14 days of incubation of the biomimetic CPP solution at 37$^{o}$C. (**E**) Graphene liquid cell without protein-mineral particles shows bubbles when exposed to a high-dose of electrons after 14 days of incubation, indicating the remaining presence of liquid in the graphene-liquid cells.
